# Conformational changes in saliva proteome guides discovery of cancer aggressiveness related markers

**DOI:** 10.1101/2023.08.04.552034

**Authors:** Daniela C. Granato, Ana Gabriela C. Normando, Carolina M. Carnielli, Luciana D. Trino, Ariane F. Busso-Lopes, Guilherme A. Câmara, Helder V. R. Filho, Romênia R. Domingues, Sami Yokoo, Bianca A. Pauletti, Fabio M. Patroni, Alan R. Santos-Silva, Márcio A. Lopes, Thaís Brandão, Ana Carolina Prado-Ribeiro, Paulo. S. L. de Oliveira, Guilherme P. Telles, Adriana F. Paes Leme

## Abstract

Diverse proteomics-based strategies have been applied to saliva to quantitatively identify diagnostic and prognostic targets for oral cancer. Considering that these potential diagnostic and prognostic factors may be regulated by events that do not imply variation in protein abundance levels, we investigated the hypothesis that changes in protein conformation can be associated with diagnosis and prognosis, revealing biological processes and novel targets of clinical relevance. For this, we employed limited proteolysis-mass spectrometry in saliva samples to explore structural alterations, comparing the proteome of healthy control and oral squamous cell carcinoma (OSCC) patients, with and without lymph node metastasis. Fifty-one proteins with potential structural rearrangements were associated with clinical patient features. Post-translational modifications, such as glycosylation, disulfide bond, and phosphorylation, were also investigated in our data using different search engines and *in silico* analysis indicating that they might contribute to structural rearrangements of the potential diagnostic and prognostic markers here identified. Altogether, this powerful approach allows for a deep investigation of complex biofluids, such as saliva, advancing the search for targets for oral cancer diagnosis and prognosis.

**Graphical Abstract:** Oral cancer progression is associated with potential structural rearrangements.

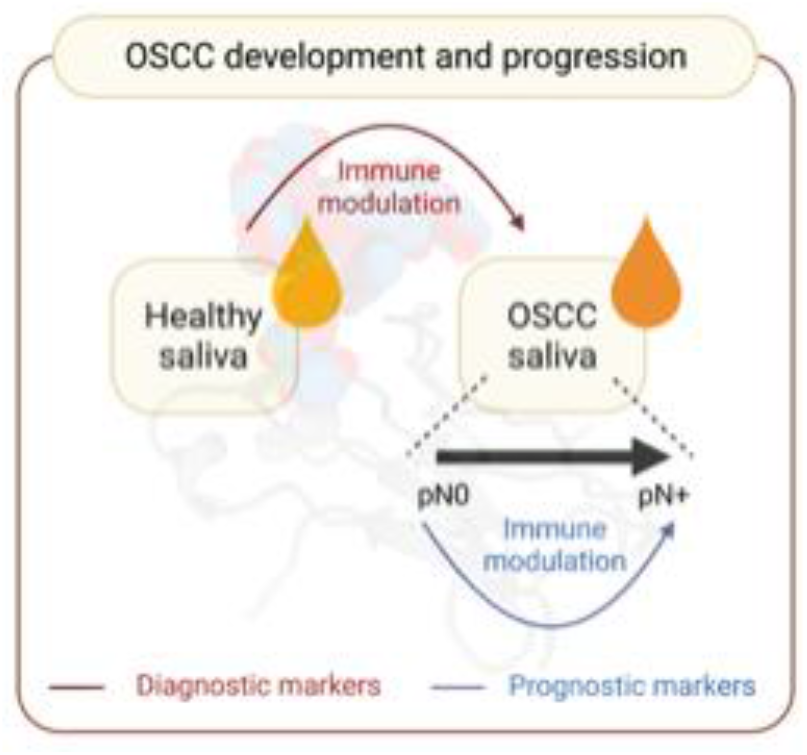

## 1. Introduction

Cancer in the oral cavity was ranked as the 16^th^ most common type of neoplasm worldwide in 2020 (Sung *et al*, 2021). It represents a health concern, with an estimation of 377,713 new cases and 177,757 deaths worldwide in 2020 (Miranda-Filho & Bray, 2020). With a five-year survival rate of around 40% to 50% (Muller & Tilakaratne, 2022), oral squamous cell carcinoma (OSCC) is the most pivotal malignancy of the oral cavity, representing more than 90% of the cases (Muller & Tilakaratne, 2022). One of the most important prognostic factors in OSCC is the presence of lymph node metastasis, and its early detection can improve the outcomes for OSCC patients, and thus in survival and quality of life. In that regard, many efforts have been made to identify prognostic biomarkers that can differentiate OSCC cases with and without lymph node metastasis (Carnielli *et al*, 2018; Chai *et al*, 2016; Chen *et al*, 2019; Kim *et al*, 2012; Sá *et al*, 2021; Schopper *et al*, 2017; Suzuki *et al*, 2016; Velmurugan *et al*, 2018; Winck *et al*, 2015; Zeng *et al*, 2016), as well as in identifying diagnostic markers capable of distinguishing OSCC cases from healthy individuals through the use of proteomics-based strategies (Sá *et al*., 2021). However, many of these proteomics studies are based on the denatured state of proteins to evaluate the abundance changes, whereas the impacts on the conformation of the proteome are still lacking information.

Considering that alterations in protein structures can affect their functional state and might be a result of events such as aggregation, misfolding, post-translational modifications (PTMs), protein-protein interactions (PPIs), or mutations, proteome-wide approaches can provide insights into these alterations and their involvement in several diseases (Mitra, 2021; Schopper *et al*., 2017). A deep structural characterization of proteins is commonly performed using X-ray crystallography, cryo-electron microscopy, nuclear magnetic resonance, and different spectroscopic methods, which have provided remarkable progress in determining complex structures (Harroun *et al*, 2022; Orellana, 2019). While most of these techniques include the use of large amounts of purified protein samples, a new mass spectrometry-based proteomic method, known as limited proteolysis-mass spectrometry (LiP-MS), has been introduced for the large-scale analysis of protein conformational changes directly from their biological matrices. In LiP-MS experiments, the use of broad-specificity proteases under native conditions dictates cleavage by the structural features of the protein (Feng *et al*, 2014). Therefore, the identification of variations in LiP-MS profiles allows the recognition of protein regions implicated in structural rearrangements (Schopper *et al*., 2017), as demonstrated in the analysis of cell lysates (Di Michele *et al*, 2015; Liu & Fitzgerald, 2016; Zuo *et al*, 2021), human plasma samples, and cerebrospinal fluid (Mackmull *et al*, 2022; Shuken *et al*, 2022; Yang *et al*, 2015), which can provide information regarding pathological conditions and guide biomarker identification (Mackmull *et al*., 2022).

Saliva is also a promising fluid for biomarker investigation because it reflects pathological and physiological alterations in local and distant parts of the body, representing a more accessible screening tool that can be repeatedly monitored over time (Muller & Tilakaratne, 2022). Moreover, the salivary proteome has been characterized in several diseases, including OSCC (Carnielli *et al*., 2018; Winck *et al*., 2015), mostly considering the traditional proteomics approach to evaluating protein abundance levels and PTMs. Saliva’s high complexity in terms of dynamic range and broad composition (Granato *et al*, 2021; Grassl *et al*, 2016) can lead to great challenges in proteome coverage and require different strategies to interrogate its proteome and detect changes associated with OSCC disease. Thus, investigating protein conformational changes in saliva can help overcome some of these challenges and contribute to the identification of novel diagnostic and prognostic markers for oral cancer.

To this end, we applied the LiP-MS strategy to saliva samples from healthy individuals compared to saliva samples from OSCC patients with or without lymph node metastasis. LiP-MS revealed 311 proteins with structural rearrangements, providing information on uniquely enriched biological processes, compared to those proteins identified and quantified based on the traditional denaturing workflow. Among the conformotypic proteins between all the conditions analyzed (healthy control and oral cancer patients, with, and without lymph node metastasis-pN0 and pN+, respectively), 51 were proteins associated with OSCC clinicopathological features. We further interrogated the source of structural rearrangements, and the results indicate potential diagnostic and prognostic proteins with various predicted PTMs, mainly disulfide bond and glycosylation, as well as proteins with glycosylation experimental evidence, which may help explain the conformational changes or the modifications that preclude enzymatic cleavage.

## 2. Results

### LiP-MS protocol development in saliva samples

For the development of this study, we followed the workflow shown in **Figure 1**. Here, we aimed to test whether protein abundance and conformational data could provide insights into functional alterations in the saliva proteome and thus identify novel diagnostic and prognostic-associated proteins based on structural changes. For this, we optimized the protocol previously developed (Cappelletti *et al*, 2021) with some modifications for (i) pooled saliva patient samples and (ii) individual patient samples. First, a pool of five saliva samples from healthy individuals was used to establish the digestion protocol for LiP treatment in saliva considering 4M urea, 4M urea with 2M thiourea, or 0.1% RapiGest. Protocol was followed with RapiGest considering the increase in sequence coverage compared to the other preparations. Next, a pooled sample of five saliva samples from pN0 and pN+ patients was equally divided and submitted for two digestion conditions: with proteinase K (+PK, *LiP*) and not treated with PK (-PK, for *tryptic digestion-only proteome, i.e., traditional proteomics workflow*). Both sample treatments were followed by standard tryptic digestion for liquid chromatography coupled with MS *in tandem* (LC-MS/MS) to identify and quantify the proteome and *LiP*-derived peptides. The results showed that pN0 vs. pN+ peptide levels upon *LiP* digestion present enriched biological processes with different profiles (**Figure S1**).

**Figure 1.**
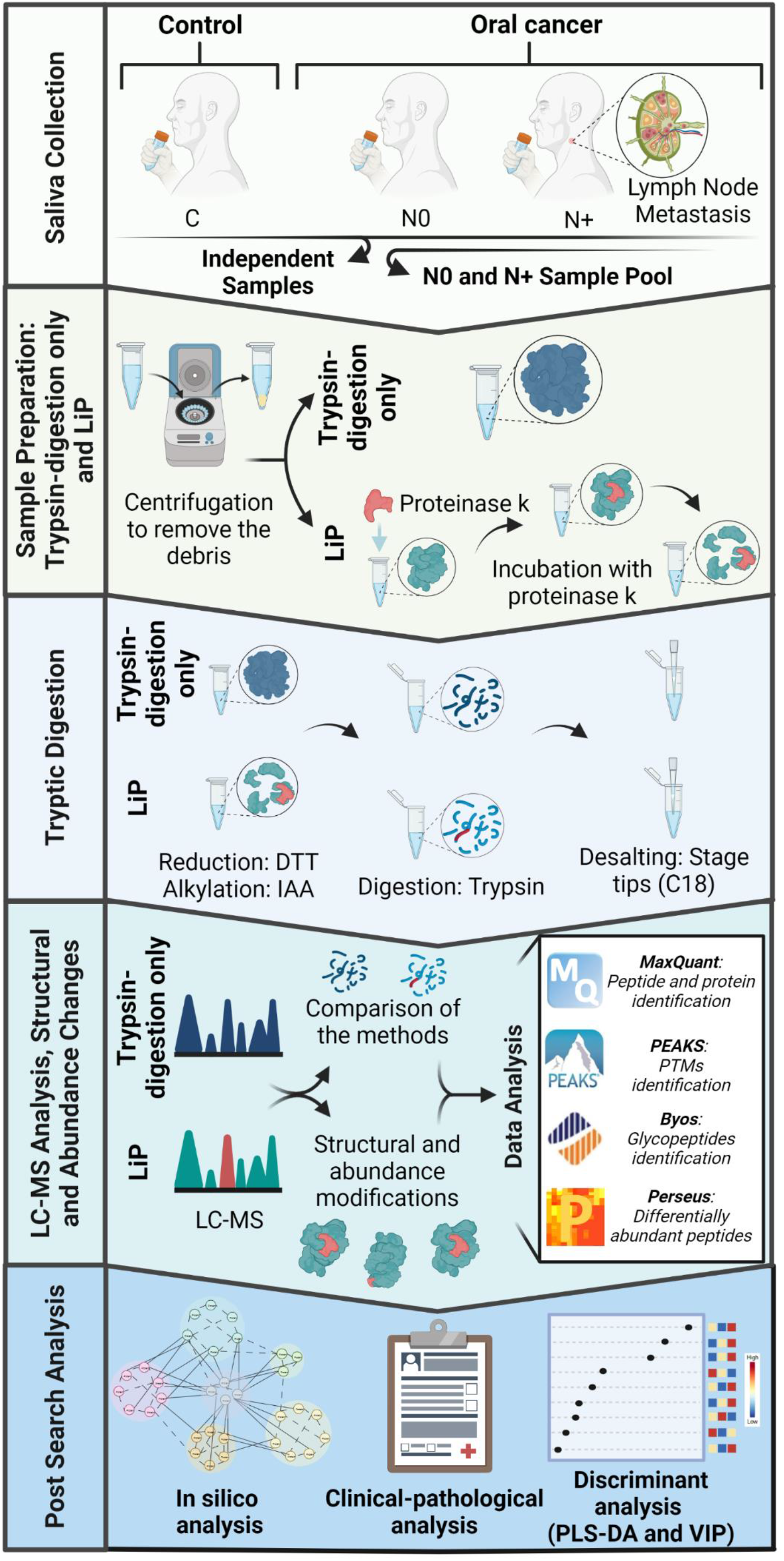
Overview of the experimental design used to map protein structural and abundance changes in the saliva of OSCC patients, with or without lymph node metastasis, in comparison to healthy patients. First, pooled saliva from the control individuals was used to optimize the digestion protocol with proteinase K (PK) treatment (LiP). Next, a pool of saliva from five individual samples from OSCC patients without lymph node metastasis (pN0) and with lymph node metastasis (pN+) were evaluated using the LiP protocol, and then the protocol was applied to independent samples from healthy individuals (n=4), and pN+ (n=4) and pN0 (n=4) patients. In LiP-MS, proteins in their native state were treated with PK. The resulting peptides’ intensities reflect changes in the conformation structures or states of proteins. Raw MS data acquired in data-dependent acquisition (DDA) mode were searched using MaxQuant and PEAKS for protein abundance (for *tryptic digestion-only proteome*) and structural changes (for *limited proteolysis*, LiP) and Byonic for the identification of glycosylated peptides. All data were analyzed by Perseus, and post-search analysis (enrichment of biological processes, dynamic protein modeling, correlation with patients’ clinical-pathological data, and ranking method analysis) was performed on significant LiP-derived peptides and proteins (ANOVA, post hoc test, p-value<0.05).

After the protocol optimization described above, we applied it to an independent cohort of saliva, including subjects from three biological groups comprised of control, OSCC- pN0, and pN+ (n=4 each group) **(Table S1–S2)**. The aims were to determine: (1) differences in protein abundance and biological processes between tryptic digestion-only proteome and LiP-MS strategies; (2) differential protein and peptides associated with OSCC diagnosis and prognosis; (3) biological functions and OSCC clinicopathological features; (4) their performance in stratifying patients by discriminant analysis; and finally, (5) diagnostic and prognostic peptides associated with PTMs.

### Comparison of sample preparation between tryptic digestion-only proteome and LiP-MS strategies

To improve protein identification/proteome coverage, samples digested only by trypsin (*tryptic digestion-only proteome)* and samples digested by PK combined with trypsin (LiP-MS) were analyzed using two different search engines: MaxQuant (database-only search) and PEAKS (database-and-*de novo* searches). The analysis of the *tryptic digestion-only proteome* resulted in the identification of 458 proteins by the database-only search, in contrast to 1,395 proteins by the database-and-*de novo* sequencing search. Likewise, 247 and 880 proteins were identified in database-only and database-and*-de novo* sequencing searches, respectively, in LiP-MS data (**Figure 2a**). Distinct dynamic range distributions were observed for both sample treatments (**Figure 2b; Tables S3–8**). The unsupervised hierarchical clustering analysis of the proteome (LiP-MS and *tryptic digestion-only proteome*) showed a partial patient grouping of pN0 and pN+ groups for all the datasets, and a slightly closer grouping of pN0 with pN+ samples in LiP-MS data for both data search strategies (**Figure 2c**).

**Figure 2.**
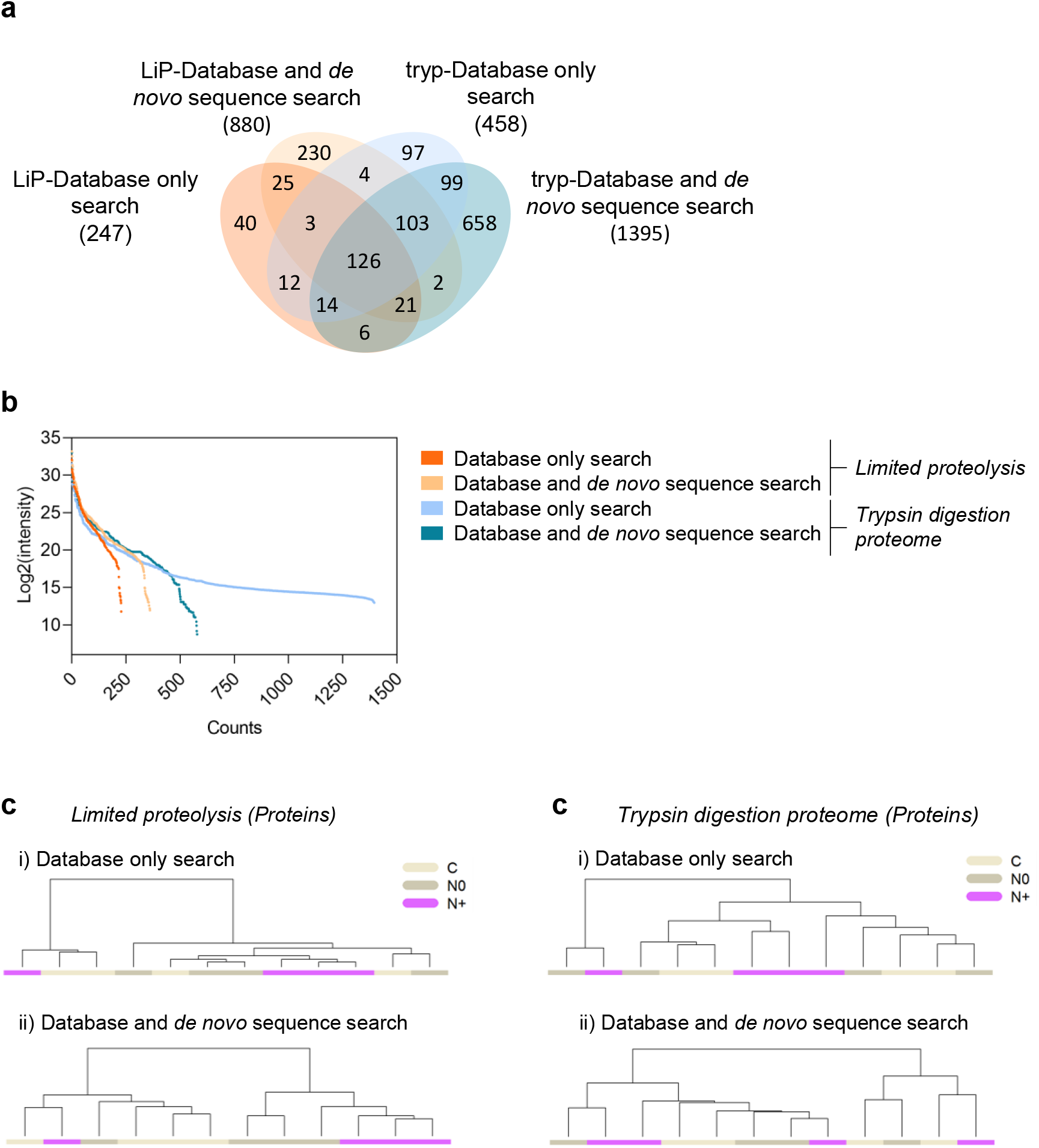
Proteome-wide scale limited proteolysis-mass spectrometry (LiP-MS) and *tryptic digestion-only proteome* analysis of OSCC saliva samples. **(a)** Venn diagram illustrating the identification overlap of proteins among the different sample treatments (tryp=trypsin digestion proteome and LiP=limited proteolysis under native conditions) using “database-only search” and/or “database-and-*de novo* sequence search”. NA: not applicable. **(b)** Dynamic range graph showing the relative distribution of protein abundance in LiP-MS and trypsin digestion-only proteome in both search strategies. **(c)** Dendrogram of the 24 LC-MS runs searched as ‘database-only’ and ‘database-and-*de novo*’ searches. Clustering analyses using Pearson correlation and Ward linkage were performed using results from proteins identified in LiP-MS and in tryptic digestion-only proteome, indicating partial grouping of biological samples (C/pN0/pN+). C=control.

### Differential peptides from LiP-MS analysis are associated with OSCC diagnosis and prognosis

We further explored the LiP-MS data to verify the association of differential peptides to patient clinical conditions, and thus gain insights into potential diagnosis (control *vs.* OSCC) and prognosis (pN+ *vs.* pN0) signatures. A total of 293 LiP peptides (from 107 proteins) identified in the database-only search presented differential abundance between pN0, pN+ groups *vs.* control group, while 1,193 peptides (from 274 proteins) identified in the database-and-*de novo* search had different levels between the three biological groups (**Figure 3a; Tables S9–10**, ANOVA, p-value<0.05, or peptides detected exclusively in one or two groups).Only 88 differential peptides from 70 proteins were commonly identified in both search strategies, indicating that these results are complementary and improve the number of identified LiP-MS peptides.

**Figure 3.**
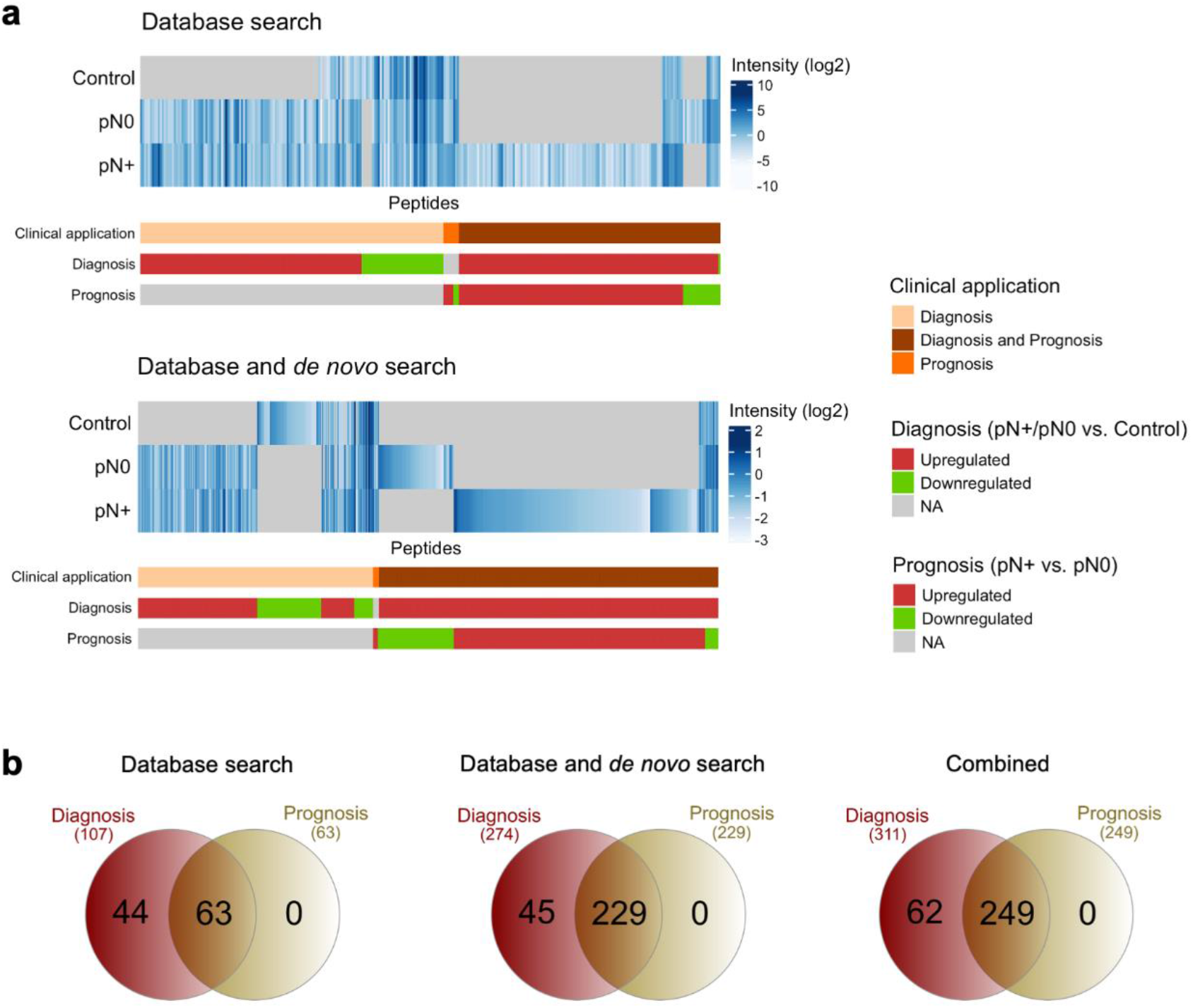
OSCC and lymph node metastasis-associated structural changes revealed by LiP-MS in the human saliva proteome. **(a)** Heatmap of peptide intensities associated with diagnosis (pN+ or pN0 vs. control; p-value ≤ 0.05; ANOVA or peptides detected exclusively in one or two groups) or prognosis (pN+ vs. pN0; p-value ≤ 0.05; ANOVA or peptides detected exclusively in one group) considering database-only search (upper panel; 293 differentially abundant peptides) or database-and-*de novo* search (bottom panel; 1,193 differentially abundant peptides). **(b)** Venn diagrams illustrating the identification overlap of source proteins of peptides associated with diagnosis and prognosis using “database-only-search” and/or “database-and-*de novo* sequence search.” NA: not applicable.

Considering data from both search analyses, most of the LiP-MS peptides (normalized by the total sum intensity of each sample) were associated with diagnosis (control vs. OSCC; 285 differential peptides from 107 proteins in the “database-based search” and 1,181 differential peptides from 274 proteins in the “database-and-de novo search” analysis; ANOVA; p-value<0.05) (**Figure 3b)**. LiP-MS peptides were also significantly associated with OSCC prognosis (pN0 vs. pN+; 140 differential peptides from 63 proteins in the “database search” and 710 differential peptides from 229 proteins in the “database-and-de novo sequencing search”, respectively, ANOVA, p-value<0.05) (**Figure 3b)**. Additionally, peptides from the two analyses were mainly upregulated (OSCC vs. control and pN+ vs. pN0) considering both diagnostic (85.2% and 85.6% of the 285 and 1,181 diagnostic peptides from “database search” and “database-and-de novo search”, respectively, ANOVA, p-value<0.05) and prognostic markers (84.2% and 74.2% of the 140 and 710 prognostic peptides from “database search” and “database and de novo search,” respectively ANOVA, p-value<0.05) (**Tables S9–10)**.

### Biological functions and OSCC clinicopathological features

Using the combined data from both searches, “database-only” and “database-and-*de novo*”, we performed a comparative analysis to retrieve biological processes or pathways enriched in the OSCC proteomes detected by structural and abundance strategies. Out of the 312 source proteins of differential and “exclusive” peptides from control vs. pN0 vs. pN+ identified in both structural (**Figure 3b**; combined Venn diagram) and abundance analysis (**Figure 3a**; ANOVA, p-value˂0.05; or peptides detected exclusively in one or two groups), 289 (92.6%) were detected exclusively by LiP-MS, and 22 (7.05%) were identified in both sample digestion approaches (**Figure 4a; Tables S9–11)**. Only one protein was detected exclusively in the abundance strategy (0.32%) (**Figure 4a; Tables S9–11)**. The 311 proteins from the LiP analysis were overrepresented in 66 GO biological processes and had potential changes in the conformation (LiP-MS analysis), including humoral immune response [false discovery rate (FDR)=3.23E-35], which is the top-1 process. Likewise, 23 proteins (differential and “exclusive” proteins from control vs. pN0 vs. pN+ of individual samples; ANOVA, p-value˂0.05) from the abundance approach (*tryptic digestion-only proteome*) were enriched for 11 significant processes, including defense response to bacterium and humoral immune response term [-log10(FDR)=5.37E-06 for both processes] (FDR ≤ 0.01) (**Figure 4b; Table S11)**. Only eight GO terms were enriched for both the structural and abundance analyses, indicating both the structural approach provided a more detailed understanding of the OSCC proteome.

**Figure 4.**
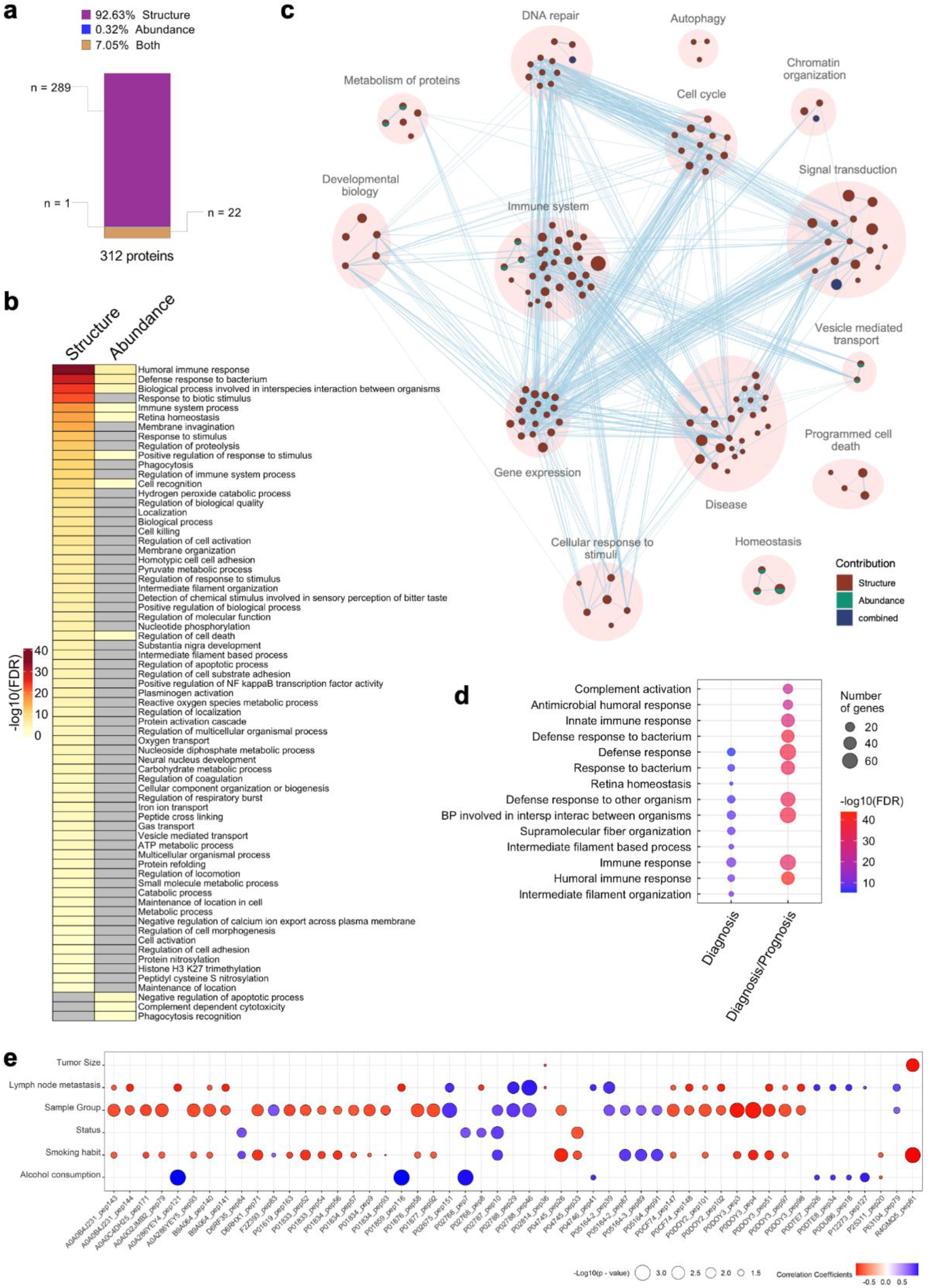
Functional analysis comparing structural (LiP-MS) and protein abundance changes in OSCC. **(a)** Number of source proteins of the differential peptides identified by structural (LiP-MS; n=311 proteins), and abundance analysis (trypsin digestion only; n=23 proteins). **(b)** Combined view of GO biological processes enriched for OSCC- associated proteins identified by LiP-MS/MS *vs.* abundance-based proteomics, shown in **(a)**. GO biological processes were enriched using the Panther tool (FDR ≤ 0.01) and overrepresented GO processes were summarized by removing redundant terms in REVIGO. **(c)** Enrichment map showing REACTOME pathways enriched for proteins containing the differential peptides identified by structural (n=311 proteins) and abundance (n=23 proteins) approaches (FDR≤0.01). Nodes represent pathways and the major functional groups are highlighted by filled circles. Enrichment significance (FDR) is represented as node sizes and pathways with common genes are shown to be connected. Nodes are colored based on supporting evidence from structural or abundance proteomes. Combined evidence was detected through data fusion and not when any of the input datasets were detected separately. **(d)** Visualization of the top-10 GO biological processes overrepresented for proteins represented in Figure 3b from peptides associated with diagnosis and prognosis (n=249 proteins) or only with diagnosis (n=62 proteins) using GSEA (FDR ≤ 0.05). **(e)** LiP-MS- generated peptides with potential clinical value for OSCC diagnosis and prognosis, indicated by their association with clinical features. Status indicates being alive or not alive. Sample group indicates control, pN0 and pN+.

In agreement with the overrepresented biological processes, enriched REACTOME pathways were also associated with immunity for source proteins of the differential peptides from structural (311 proteins) and abundance (23 proteins) analyses (FDR≤0.01) (**Figure 4c; Table S12)**. Interestingly, while 102 pathways were overrepresented in the LiP-MS combined dataset, nine pathways associated with protein metabolism, the immune system, and homeostasis were enriched in both approaches (**Figure 4c)**. Three pathways significantly associated with DNA repair, signal transduction, and chromatin organization were overrepresented only when the structural and abundance datasets were combined. Once again, this analysis demonstrated that the LiP-MS approach outperformed the abundance analysis in terms of differential protein changes and enrichment processes and pathways’, thus providing a deeper view of the OSCC saliva proteomes (**Figure 4c**).

To evaluate GO biological processes enrichment from a clinical perspective, the combined diagnostic and prognostic candidate proteins from **Figure 3b** and **Tables S9– S10** (source proteins of the differential and “exclusive” peptides from control vs. pN0 vs. pN+ of individual samples; ANOVA, p-value ˂ 0.05; or peptides detected exclusively in one or two groups), comprising 62 proteins from the diagnostic peptides (differential abundance in control vs. OSCC) and 249 proteins from peptides with both diagnostic (differential abundance in control vs. OSCC) and prognostic (differential abundance in pN0 vs. pN+) values, identified by “database” and “database-and-*de novo*” searches, were evaluated (**Figure 4d**). Immune processes, mainly humoral responses, were overrepresented when diagnosis and prognosis candidates were combined. Even though there are only a few distinct enriched biological processes, there is a notable increase in gene number representation in innate immune-related processes (complement activation and antimicrobial humoral response) (**Figure 4d; Table S13)**.

We then investigated whether the differential diagnostic and prognostic LiP candidate peptides were associated with clinical data, revealing 249 potential conformotypic peptides with a significant association with prognostic features (linear regression p-value <0.05, and Pearson’s R-squared>0.5). By applying a stringent criterion (correlation with ≥2 clinicopathological features), a subset of 51 peptides correlated with alcohol and smoking consumption, sample group, lymph node metastasis and tumor size (**Figure 4e; Table S14**).

### LiP-peptide performance in discriminating patients

We investigated the ability of differential protein-derived peptides (control vs. pN0 vs. pN+, ANOVA, p-value <0.05) identified in the LiP-MS strategy derived from “database-and-*de novo*” searches in distinguishing the sample groups using different ranking methods. As a result, the top-five peptides derived from TKT1, IGLC2, IGLL5, TKT2, and IGHA2 proteins had the best ranking performance considering the sample groups. We selected transketolase (TKT-1 and TKT-2) among the top-five targets based on its common evidence among the different ranking strategies (**Figure S2**) and its potential prognostic value when associated with clinicopathological features relevant for OSCC, such as lymph node metastasis (pN0 vs. pN+). TKT was also selected as a target for further analysis because it was identified in OSCC tumor tissue as differentially abundant between non-malignant cells from primary tumors compared to lymph nodes (upregulated in lymph nodes, q ≤ 0.05; Student’s t-test followed by Benjamini–Hochberg correction), as previously reported by our group (Busso-Lopes *et al*, 2022).

To explore the potential conformational changes in TKT (TKT-1 and TKT-2), we visualized the TKT conformotypic peptides on the tridimensional structure (PDB ID 3MOS) using Chimera, indicating the four conformotypic peptides present in each monomer, which are exclusive or upregulated in pN+, indicating conformational changes associated with lymph node metastasis (**Figure 5a**). The peptides 1 and 3 present amino acids that are involved in thiamine binding while the other two are annotated in the Uniprot database with N6-acetyllysine in 538 and phosphoserine in 287 and 295 (**Figure 5a**), which may explain the structural changes between the healthy and diseased states.

**Figure 5.**
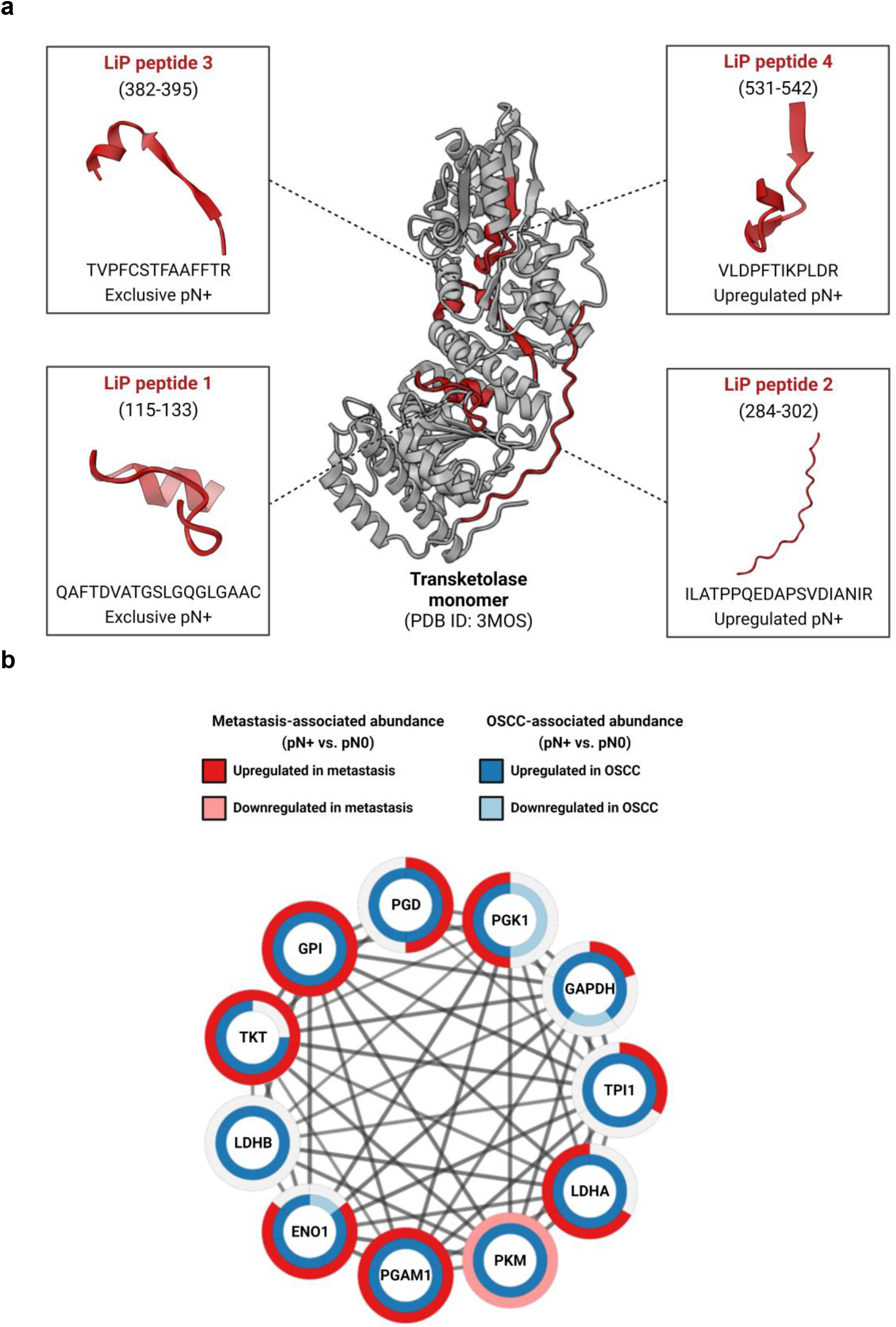
TKT is a strong candidate for conformational changes associated with OSCC redox and metabolic processes. **a)** Conformotypic peptides are highlighted in red. **b)** The STRING network of TKT and functional associated proteins that the LiP-MS strategy indicates structural changes. Each node represents a protein, and edges connecting nodes represent high confidence functional associations (STRING score >0.7).

TKT also shows functional associations with other proteins (LDHA, LDHB, PKM, ENO1, GPI, GAPDH, PGK1, PGAM1, and PGD) identified in this LiP-MS strategy using STRING (**Tables S9–10**), revealing a network proteome that could also undergo structural changes associated with oral cancer (**Figure 5b**).

### Diagnostic and prognostic peptides associated with PTMs

Considering that phosphorylation can modify the structure of prognostic TKT protein, as demonstrated above, we further used the UniProt PTM database to verify predicted PTMs in proteins and peptides from the diagnostic or prognostic groups revealed by LiP-MS, as shown in **Figure 3a** (**Tables S15–21**). To evaluate the role of PTMs distant to the LiP peptide and in the peptide sequence, we opted to evaluate the representation of PTMs considering the full sequence of the protein and the peptide sequence or vicinity (+/- 10 amino acids), respectively.

For the diagnostic proteins, the most frequent PTM was the disulfide bond (23.1%), followed by glycosylation (15.1%) and phosphoserine (11.9%), as shown in **Figure 6a**. Indeed, these were also the three most predominant PTMs for the prognostic (**Figure 6b**) group, with glycosylation as the top hit. In terms of peptide sequences identified in the LiP-MS, the diagnostic (**Figure 6c**) and prognostic (**Figure 6d**) associated-LiP peptides indicated that the highly dominant PTM was disulfide bond, contributing 74% and 81.8%, respectively. We can also observe the potential contribution of other modifications, such as phosphorylation and glycosylation. Although these LiP-MS peptides contained predicted PTMs, they were not identified with their respective PTM counterparts by LC-MSMS, although they may still have them.

**Figure 6.**
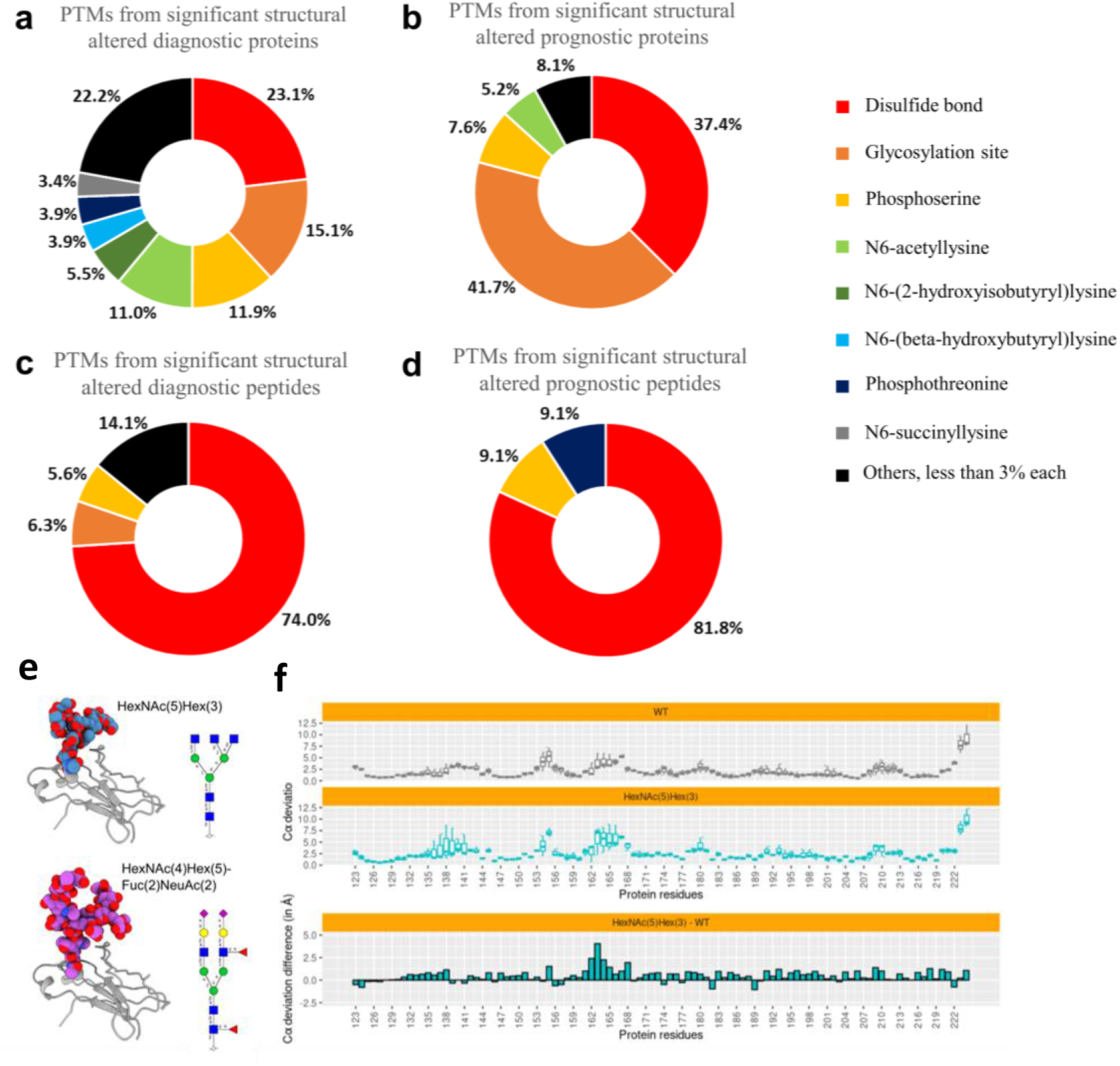
Distribution of PTMs in LiP differential proteins in saliva. The distribution of PTMs in a significant LiP peptide sequence or vicinity (±10 amino acids) or proteins was analyzed considering the following groups: **(a)** diagnostic proteins, **(b)** prognostic proteins, **(c)** diagnostic peptides, and **(d)** prognostic peptides. The three most predicted PTMs are disulfide bonds, glycosylation sites, and phosphoserine. Moreover, disulfide bonds are the major PTM when comparing the significantly altered peptides. (**e**) 3D models of the glycans HexNAc(5) Hex(3) (in blue, upper model) and HexNAc(4)Hex(5)Fuc(2)NeuAc(2) (in purple, bottom model) bound to the IGHA1 Fc domain through the Asn144 residue. A schematic view of each glycan chain is also presented. (**f**) Analysis of 100 ns (in triplicate) molecular dynamics simulations in the absence (WT) or in the presence of the HexNAc(5) Hex(3) glycan. The two upper plots show the distance (only considering Cα atoms) between each IGHA1 residue in the simulation trajectory and in a reference initial structure. The reference structure corresponds to the first frame collected from the WT simulation. Higher values mean that the residue deviates from its position in the reference structure during the trajectory. Each boxplot presents the meaning of deviations computed along the simulation trajectory in triplicate. The bottom plot shows the deviation difference per residue upon subtraction of HexNAc(5) Hex(3) glycan distances by WT distances. Higher values mean that a higher deviation was observed in the simulation with IGHA1 bound to HexNAc(5) Hex(3) in comparison to the simulations without glycan.

Considering that glycosylation showed the highest contribution (41.7%) among all the PTMs in the prognostic protein group, we further explored our LiP proteomic data using Byonic node (Protein Metrics Inc., CA, USA) within Proteome Discoverer v.2.4.0.305 (Thermo Scientific) for the identification of glycopeptides (**Table S22**). Results indicated 34 distinct glycopeptides (control vs. pN0 vs. pN+ ANOVA, not differential) and 92 non-glycosylated differential peptides (control vs. pN0 vs. pN+ ANOVA, p-value <0.05) (**Table S23–25**).

To provide further insights into how glycosylation could produce differential peptide profiles, we selected the IGHA1 Fc domain, which has a known 3D structure (PDB ID: 6UE7), for *in silico* molecular modeling. We modeled each of the two identified glycans, (HexNAc[5] Hex[3] and the HexNAc[4]Hex[5]Fuc[2]NeuAc2)) (**Figure 6e**), attached to the IGHA1 Asn144 residue and then performed molecular dynamics (MD) simulations to assess how the glycans could affect the IGHA1 structure, peptides identified in the control only and not in the other groups (pN0 and pN+). As expected, the

MD simulations evidenced the high flexibility of both glycan chains in the context of the IGHA1 chain (**Figure S3**). By computing the Cα position deviation of each IGHA1 residue over the simulation trajectory in relation to an initial reference structure, we observed a higher deviation in several residues when IGHA1 was bound to glycan chains compared to simulations without glycans (WT) (**Figure 6f and Figure S3**). This finding may indicate that the presence of glycans could affect the overall IGHA1 structure and, therefore, alter the proteolytic interaction with the proteinase K and, ultimately, contribute to the conformational changes detected by LiP-MS analysis. Additionally, the flexibility of the glycans observed in the simulations indicates that the glycans can cover an extensive area of the IGHA1 surface, which includes putative contacts with identified conformotypic peptides, which are conformation-specific peptides produced during protease digestion that can be used as structural probes (**Figure S3**). Thus, the presence of glycans alone could change the structural interface targeted with the proteinase K and impair the accessibility of this enzyme to IGHA1 proteolytic target sites.

## 3. Discussion

Classical experiments evaluating the proteome of saliva, the fluid in which the primary oral tumor is embedded, have led to the identification of promising proteins as potential diagnostic and prognostic signatures for OSCC, improving our understanding, and potential management of the disease (Carnielli *et al*., 2018; Granato *et al*., 2021; Winck *et al*., 2015). However, conformational changes in proteins present in saliva as potential biomarkers have not been previously investigated in oral cancer. In this scenario, saliva has a great advantage for structural analysis since it does not require protein extraction under denaturing conditions; therefore, its native proteome can be directly studied by LiP-MS workflow (Cappelletti *et al*., 2021; Leuenberger *et al*, 2017).

To date, LiP-MS is considered one of the main MS approaches enabling the probing of proteome structural alterations, providing information on both protein abundance under denaturing conditions (herein called *tryptic digestion-only proteome*) and protein structural changes (LiP-MS strategy). This approach uses the rate of enzyme-driven non-tryptic proteolytic cleavage to infer changes in protein structure, typically associated with changes in function that can occur for many reasons, such as mutation, aggregation, folding, proteoforms, stability, molecular binding, PTMs or changes in the properties of the matrix (Merl-Pham *et al*, 2019). Regulation of cellular processes, such as signaling cascades, might rely on these types of events rather than on changes in the protein abundance denatured state.

Through LiP-MS combined with classical proteome workflow, we investigated changes in the abundance and structure of proteins from the saliva of OSCC patients and from control individuals (**Figure 1**), aiming to identify conformational changes with clinical value. For this, we combined the output of three different search algorithms (MaxQuant, PEAKS, and Byonic) to generate a list of conformotypic peptides that is, peptides with conformational changes identified by LiP-MS. Our results show that tryptic digestion of the saliva proteome reflects higher intragroup heterogeneity, probably because the proteome under denatured conditions is more readily available for cleavage, when compared to LiP-MS treatment, which has a more uniform distribution intragroup (**Figure 2**). Notwithstanding, clustering analysis of LiP peptide data enabled a slightly better grouping between the control and OSCC samples (**Figure 2**). However, clustering analysis was still unable to segregate OSCC samples based on lymph node status, as observed in other head and neck squamous cell carcinoma (HNSCC) proteomes (Busso-Lopes *et al*., 2022; Carnielli *et al*., 2018). This must be related to the clinical and biological variability of OSCCs once tumors at the same clinical stage may have distinct clinical outcomes (Fisher *et al*, 2013). Despite this, a total of 293 and 1,193 differential peptides (ANOVA, p-value<0.05) were identified in LiP-MS data with MaxQuant and PEAKS tools, respectively, comparing pN0, pN+, and control groups. This result indicates potential diagnostic and prognostic candidates (**Figure 3**).

Our study provides complementary information on the 311 proteins structurally altered in the saliva proteome of OSCC patients when combining different search algorithms (**Figure 4a**). Despite the LiP-MS strategy indicating a lower dynamic range of the proteome in saliva samples (**Figure 2**), these results pointed to more enriched biological process networks when compared to the traditional proteomics workflow results (**Figure 4b–c**). Moreover, to explore the potential clinical relevance of OSCC on LiP-MS hits (**Figure 4e**), we investigated the association of peptide abundance with clinicopathological features. This analysis showed an association between 51 peptide abundances with OSCC and lymph node metastasis, revealing potential diagnosis and prognosis markers, respectively (**Figure 4e**).

To further explore the clinical value of LiP-MS data, the ability to distinguish the OSCC patients was tested, and transketolase (TKT), an enzyme that participates in the pentose phosphate pathway, was identified as a potential structural marker in this study. TKT was upregulated in the pN+ condition (from database-and-de novo search analysis) and showed high performance in discriminating patient conditions by different ranking method analysis (**Figure 5**). This protein was also previously identified with higher abundance in non-malignant cells from lymph nodes compared to primary tumor’s cells (Busso-Lopes *et al*., 2022). Although TKT has a recognized prognostic role in tumoral tissue, its contribution in saliva has yet to be determined. The association between proteins that are involved in both OSCC tissue and saliva has been reported in a previous work from our group (Carnielli *et al*., 2018) which demonstrated that a prognostic signature from tumor regions has similar levels in the saliva of OSCC patients.

TKT enzymes are phosphorylated in Thr380 by AKT (Saha *et al*, 2014), and the structural changes detected by LiP-MS encompass this region, implying a role of phosphorylation in TKT conformational change, more specifically in metastasis condition (Xu *et al*, 2016). However, no phosphorylated site was identified in this protein by “dataset-and-*de novo* search” data. TKT is functional in its dimeric form, and it is known to bind to the cofactor thiamine pyrophosphate (or diphosphate), which is required for TKT to catalyze its reaction. The binding to TKT occurs through hydrogen bonds and hydrophobic interactions in several residues (Mitschke *et al*, 2010). Binding to thiamine does not induce conformational changes in the protein (Deshpande *et al*, 2019) since it contains a loop that is open or closed when bound or unbound to thiamine, respectively. However, binding to thiamine can modify the accessibility to proteinase K in this region, generating conformotypic peptides.

Considering the large evidence of disulfide bonds and glycosylation in previously described saliva traditional proteomics data (Grassl *et al*., 2016), we investigated annotated PTMs in LiP-MS results that can guide the observed conformational changes. Glycosylated residues and disulfide bonds were the most annotated modifications in significant LiP-MS proteins, while differential LiP-MS peptide sequences were mostly annotated with disulfide bond and phosphorylation sites (**Figure 6c**). Although it explains that the PTM functional role in salivary proteins is beyond the scope of this study, it is noticeable that the LiP-MS strategy can retrieve protein/peptide with clinical relevance, enriching disulfide bond modification. As proof of concept, we investigated a highly predicted PTM in our LiP-MS data (**Figure 6e-f**). Although glycopeptides are often present in a lower dynamic range and exhibit reduced ionization efficiency in mass spectrometry compared to non-glycosylated peptides (Oliveira *et al*, 2021), we were able to identify *N*-glycosylated peptides in our non-enriched samples data, including *N*-glycans from the main classes: highly truncated, paucimannosidic, oligomannosidic and complex/hybrid *N*-glycans (**Table S22**). To explore whether the identified glycosylation is associated with conformational changes, we evaluated the IGHA1 protein, in which glycopeptides were identified in the glycopeptide search only in the control samples (**Figure 6b-c**). In our data, two different IGHA1 glycopeptides, one carrying HexNAc (5) Hex (3) and the other carrying HexNAc(4) Hex(5)Fuc(2)NeuAc(2) *N*-glycan, were identified. *In silico* modeling suggests a change in conformational upon the presence of glycans, which can also be attributed to a change in the accessibility of the enzyme, although the respective non-glycosylated peptides were not identified in the other groups, with protein coverage of about 90% in the control, 91% in pN0 and 89% in pN+.

In summary, this conformational comprehensive approach using saliva revealed that the structural readout captured more altered proteins and biological processes, suggesting that i) the integration of these approaches with classical proteomics workflow can improve the detection of novel and robust diagnostic and prognostic-associated proteins, mainly in high protein dynamic range matrix; ii) can be a powerful strategy to detect altered biological processes assisting in the search for molecular events underlying physiological and pathological events; and iii) allows the prioritization of novel proteins and biological processes associated with OSCC and not revealed by traditional proteomics workflow methods, selecting potential candidate as markers for monitoring in the path of disease or therapeutic targets. Altogether, our findings show that LiP-MS can identify structural changes in saliva associated with aggressiveness and with relevance to the diagnosis and prognosis of the disease. This study is a pioneer in the field, showing a deeper structural salivary proteome, while opening new possibilities for studies.

## 4. Materials and Methods

### Patient selection

This study comprised two patient cohorts: the first encompassed a pool of saliva samples from five patients with lymph node metastasis (pN+; “p” indicates the presence of metastasis in the lymph node histologically confirmed) and without lymph node metastasis (pN0) for protocol optimization, and an independent patient cohort including healthy control (n=4) and OSCC patients (n=8, N0=4, N+=4). Only patients who had active oral malignant lesions at the time of saliva collection, in the presence or absence of lymph node metastasis (pN+ and pN0, respectively) were collected. The detailed clinical and pathological information for the eight OSCC patients and the clinical information of the four healthy individuals enrolled in this study are summarized in **Table S1**.

This study was approved by the Ethics Committee of the Instituto do Câncer do Estado de São Paulo (ICESP), São Paulo, SP, Brazil, through protocol CAAE 30658014.1.1001.0065, and by the Ethics Committee of the Piracicaba Dental School (FOP), São Paulo, SP, Brazil, through protocol CAAE 51762415.5.0000.5418. All participants provided written informed consent. The study was conducted following the Declaration of Helsinki and was performed following the Strengthening the Reporting of Observational Studies in Epidemiology (STROBE) statement (von Elm *et al*, 2008).The procedures used for saliva sampling ^15^ and annotation were performed following the guidelines and experimental protocols approved by both ethics’ committees.

### Saliva collection

All saliva samples were collected from individuals who were advised not to eat for at least 1 h, then rinsed their mouths with 5 mL of drinking water, and saliva was subsequently harvested without stimulation into a sterile plastic bottle for 5 min^15^. Frozen stocks of saliva were stored at −80°C until further used^15^. Before use, saliva cellular debris was first harvested by centrifugation at 1.500 x g for 5 min at 4^°^C and the supernatant was collected and transferred to a fresh micro tube. Protein concentration was determined with the Bradford assay (Bradford Assay Kit, Thermo Fisher Scientific).

### Saliva preparation through limited proteolysis (LiP)

For protocol optimization, three distinct protocols with different denaturing agents and concentrations (4M urea, 4M urea, and 2M thiourea, or 0.1% RapiGest) were tested in a pool of healthy individual’s saliva samples for protein digestion following LiP-MS. The best protocol (RapiGest) was selected based on the highest number of proteins and peptides identification. Ten micrograms of each proteome extract were subjected to tryptic digestion (as described below) for protein abundance change evaluation (*tryptic digestion-only proteome*) and to access protein structural changes. Another 10 μg was submitted for the LiP-MS approach, which included a double-protease digestion step with a nonspecific protease followed by digestion with trypsin. First, for the LiP approach, proteinase K (PK) from *Tritirachium album* (Sigma Aldrich) was added to the samples under native conditions at an enzyme/substrate (E:S) ratio of 1:100 (w/w) and incubated for 1 min at 25°C with agitation^17^. In parallel, the same volume of water was added to the *tryptic digestion-only proteome* samples and incubated for 1 min at 25°C with agitation. PK digestion was stopped by heating samples for 5 min at 98°C in a thermocycler, followed by the addition of RapiGest (Waters) to a final concentration of 0.1% for protein denaturation. The same procedure was applied to *tryptic digestion-only proteome* samples. Both the LiP-MS and *tryptic digestion-only proteome* samples were then subjected to complete tryptic digestion in denaturing conditions, as described below.

### Tryptic digestion

The saliva samples analyzed in this study were reduced by incubation with dithiothreitol (DTT) (Sigma Aldrich) to a final concentration of 5 mM for 25 min at 56 °C. Next, the alkylation of free cysteine residues was achieved by adding iodoacetamide- IAA (Sigma Aldrich) to a final concentration of 14 mM for 30 min at room temperature in the dark. Samples were digested with sequencing-grade porcine trypsin (Promega) at an enzyme/substrate ratio of 1:100, added twice to the samples after 4 h at 37°C, and incubated for 16 h at 37°C. Protease digestion was quenched by lowering the reaction pH (< 3) with a final concentration of 0.5% trifluoroacetic acid. The peptide mixtures were loaded onto stage-tip C18 for sample desalting and eluted with 80% acetonitrile in 0.1% formic acid. After elution, the peptides were dried in a vacuum centrifuge, re-solubilized in 0.1% formic acid, and analyzed by mass spectrometry. The run order was randomized using an R (v3.4.0) environment to prevent systematic bias during MS acquisition (**Table S2**).

### LC-MS/MS data acquisition

Peptide digests (2μg) from LiP-MS and *tryptic digestion-only proteome* samples were analyzed on an LTQ Orbitrap Velos (Thermo Fisher Scientific, San Jose, CA) mass spectrometer coupled to a nanoflow liquid chromatography on an EASYnLC system (Proxeon Biosystems, Odense, DK) with a Proxeon nanoelectrospray ion source. Peptides were subsequently separated in a 2%–90% acetonitrile gradient in 0.1% formic acid using a PicoFrit analytical column (20 cm × ID75, 5 μm particle size, New Objective) at a flow rate of 300 nL/min over 220 min, in which a gradient of 35% acetonitrile was reached in 173 min. The nanoelectrospray voltage was set to 2.2 kV, and the source temperature was set to 275 °C. The instrument methods employed for LTQ Orbitrap Velos were set up in data-dependent acquisition (DDA) mode. Full-scan MS spectra (m/z 300–1,600) were acquired in the Orbitrap analyzer after accumulation to a target value of 1E^6^. Resolution in the Orbitrap was set to r = 60,000, and the 20 most intense peptide ions (top 20) with charge states ≥1 were sequentially isolated to a target value of 5,000 and fragmented in the high-pressure linear ion trap by collision-induced dissociation (CID) with a normalized collision energy of 35%. Dynamic exclusion was enabled with an exclusion list size of 500 peptides, an exclusion duration of 60 seconds, and a repetition count of 1. An activation Q of 0.25 and an activation time of 10 ms were used. A complete overview of the experimental workflow is given in **Figure 1**.

### Peptide and protein identification

Raw files from both digestion approaches were searched against the *Human* Uniprot database (version June 2018, 93,599 protein sequences, 36,574,184 residues) using the MaxQuant search engine (version 1.5.8) and PEAKS Studio X (version 7.5, Bioinformatics Solutions Inc). For the *tryptic digestion-only* proteome on MaxQuant, up to two missed cleavages were allowed, and the cleavage of KP and RP peptide bonds was excluded (**Table S3**). For the MaxQuant search of LiP-MS samples, a semi-specific tryptic digestion was set (**Tables S5–6**), while for *tryptic digestion-only proteome* samples, a fully tryptic digestion was applied (**Table S3**). Cysteine carbamidomethylation (+57.0214 Da) and methionine oxidation (+15.99492) were allowed as fixed and variable modifications, respectively; monoisotopic peptide tolerance was set to 10 ppm, and fragment mass tolerance was set to 1 Da for both digestion strategies. The FDR was estimated using the target-decoy method (Choi *et al*, 2008) and set to 1% at the peptide and protein levels.

For PEAKS analysis, the database combined with *de novo* searches were performed. In the latter case, PTM exploratory search was also included in both types of samples (*tryptic digestion-only proteome* and *LiP)*, and all 313 variable modifications present in the PEAKS PTM database were defined and used as variable modifications. Also, a fully tryptic digestion rule was used for protein abundance quantification for the *tryptic digestion-only proteome* (**Table S4**), while using the semi-specific tryptic rule for LiP-MS samples (**Tables S7–8**). The FDR was estimated using the target-decoy method^31^ and set to 1% at the peptide and protein levels.

Also, Byonic node (Protein Metrics Inc, CA, USA) within Proteome Discoverer v 2.4.0.305 (Thermo Scientific) for the identification of intact glycopeptides in both types of samples (*tryptic digestion-only proteome* and *LiP*), using mammalian glycan database (containing 134 distinct glycans provided by the software) for glycan composition information and Uniprot database (version June 2018, 93,599 protein sequences, 36,574,184 residues) for protein search. For LiP-MS, a semi-specific tryptic rule was used for the LiP-MS samples (**Table S22**). All searches were filtered to <1% FDR at the protein level and 0% at the peptide level by using a protein decoy database (*37*) (**Table S22**). Only *N*-glycopeptides confidently identified with PEP 2D scores < 0.001 were considered (*28*) (**Table S22**).

### LiP-MS data analysis

In the LiP-MS samples, only fully- and semi-tryptic peptides that had at least one identified peptide were used for quantification (**Tables S5**, **S6–8, and S22**), while in the *tryptic digestion-only proteome* samples only fully tryptic proteotypic peptides were used for protein abundance analysis (**Tables S3–4**).

LiP-MS data obtained by all search engines (MaxQuant, PEAKS, and Byonic) were individually normalized at the peptide level by dividing each peptide intensity by the sum intensity value of each sample (**Tables S5–6, S8, and S23**). Normalized peak areas were used for quantitation in LiP-MS samples, considering fully and semi-tryptic peptides. Group comparison analysis of peptides (LiP-MS samples) and proteins (*tryptic digestion-only proteome*) was performed independently within Perseus (version 1.6.5) for all search algorithms (**Tables S9–10 and Tables S22–23, respectively**). At the protein level, LFQ values transformed in Log2 were used for quantification.

### Definition and annotation of prognostic and diagnostic markers

Normalized intensity values for LiP-MS were performed at the peptide level from MaxQuant, and PEAKS searches were used to determine differentially protein conformation-based changes among control, pN0, and pN+ conditions in Perseus v. 1.3.0.4 software (ANOVA followed by Tukey’s test, p-value<0.05) (Tyanova *et al*, 2016). Glycopeptides identified in the Byonic search were used exclusively for qualitative analysis to access the role of glycans on *in silico* protein conformation, once none of the glycosylated peptides were statistically significant (ANOVA, p-value>0.05) considering the selected criteria. Peptides with significant differential abundances or exclusively detected in one or two out of three analyzed conditions were grouped as diagnostic or prognostic markers considering their association with OSCC, following the criteria described below (**Tables S9–10**):

i. Potential diagnostic markers: peptides differentially abundant between control and OSCC (pN0 and/or pN+, ANOVA, p-value<0.05). This category also
ii. peptides detected “exclusively” in control or OSCC conditions. Diagnostic markers with higher abundance in OSCC (pN+ and/or pN0) compared to the control or those exclusively detected in OSCC were termed “upregulated,” while peptides exhibiting lower expression in OSCC (pN+ and/or pN0) compared to the control or those exclusively detected in the control group were named “downregulated.”
iii. Potential prognostic markers: peptides differentially abundant between pN+ and pN0 (ANOVA, p-value ≤ 0.05) and peptides exclusively identified in N+ or N0 conditions. Prognostic markers exhibiting higher expression in pN+ compared to pN0 or those exclusively detected in pN+ were termed “upregulated”, while peptides exhibiting lower expression in pN+ compared to pN0 or those exclusively detected in pN0 were termed “downregulated”.

Common identifications between prognostic and diagnostic proteins were visualized in Venn diagrams using the InteractiVenn tool (Heberle *et al*, 2015).

### Biological process enrichment analysis

For the potential diagnostic and prognostic markers, GO biological processes were enriched using the Molecular Signatures Database from Gene Set Enrichment Analysis software (Fisher’s Exact Test, p-value <0.05) (Liberzon *et al*, 2011; Subramanian *et al*, 2005), and the top-10 overrepresented GO terms were simultaneously visualized for proteins with potential diagnostic and prognostic value, combined with source proteins from diagnostic peptides.

We also tested the proteins with significant abundance changes in *tryptic digestion-only proteome* and the proteins with significant conformational based changes for LiP-MS proteome samples among the control, pN0, and pN+ groups. Overrepresented GO biological processes were determined with Panther (Mi *et al*, 2021; Thomas *et al*, 2022) tool (FDR ≤ 0.01; Bonferroni correction) and summarized by removing redundant terms in REVIGO (Supek *et al*, 2011). Molecular pathways were enriched using the ActivePathways (Paczkowska *et al*, 2020) tool with hypergeometric tests to determine overrepresented REACTOME terms (FDR ≤ 0.01). These terms were used as input for the EnrichmentMap (Merico *et al*, 2010; Reimand *et al*, 2019) plugin in Cytoscape v.3.9.1 (Cline *et al*, 2007) for network visualization and their coloring according to supporting evidence. Enrichment maps for pathways overrepresented in structural and abundance analysis were visualized with a Jaccard and overlap combined coefficient of 0.6. Major biological themes and their relationships were represented as clusters of similar pathways. A p-value threshold of 0.05 was set to determine significance (**Tables S11–13**).

### Feature analysis

To explore which features were enriched among the significantly altered *LiP* peptides, we used Typic (Pauletti *et al*, 2023) to retrieve Uniprot database features (eg. amino acid modifications, PTMs including glycosylation and disulfide bond, mutations) with experimental evidence from potential diagnostic and/or prognostic LiP markers (**Tables S15–21**).

### Prioritization of conformotypic peptides associated with OSCC diagnosis and prognosis

Prioritization was based on the following criteria: 1) top hit in ranking attribute method analysis; 2) clinical correlation with prognostic features; 3) identifications in previous studies as differentially abundant proteins from non-malignant cells of primary tumors compared to committed lymph nodes (Busso-Lopes *et al*., 2022).

Three filter approaches implemented in the data mining tool Waikato Environment for Knowledge Analysis (WEKA) v.3.8.5 were applied in LiP data set. They were i) information gain attribute evaluation, ii) chi-square attribute evaluation, and iii) SVM attribute evaluation. Also, the ranking method of search, german_credit, is applied to the dataset.

To investigate the diagnostic or prognostic value of the significant conformotypic peptides, a linear regression analysis was performed using R code to evaluate the linear relationship between peptide abundance and clinicopathological variables. P-value <0.05 was used to define significance (**Table S14**). The Pearson-product moment correlation coefficient was calculated to infer the strength of the association between the described variables. Only the peptides that corresponded to peptides associated with patient clinicopathological data (alcohol consumption, smoking habit, sample group, status, lymph node metastasis, and tumor size) with prognostic value were further investigated. Also, peptides were evaluated regarding their distinguishing performance in ranking score.

To visualize the prioritized target, TKT, and its conformational peptides in a 3D view, the prioritized protein was analyzed in Chimera X using pdb data.

### Molecular dynamic simulation

To model the N-glycosylation of the immunoglobulin heavy constant alpha 1 (IGHA1), the IGHA1 3D structure was obtained from the Protein Data Bank (PDB ID: 6UE7) and was preprocessed to keep only the Ig-like 2 (UNIPROT ID: P01876) domain which is the target of N-glycosylation at position 144. The N-glycan residues (HexNAc[4] Hex[5]Fuc[2]NeuAc[2] and HexNAc[5]Hex[3]) were modeled using the CHARMM-GUI (Jo *et al*, 2008) and Glycan Modeler module (Park *et al*, 2019). To access the dynamics of the glycans and their impact on the IGHA1 structure, we used three initial models: (1) the IGHA1 Ig-like 2 domain without glycan, (2) with the HexNAc(4) Hex(5)Fuc(2)NeuAc(2) N-glycan, or (3) with the HexNAc(5)Hex(3) N-glycan. For MD simulations, the N-terminal and C-terminal caps were added using CHARMM-GUI and the Solution Builder module was used to generate a rectangular solvation box (edge distance of 10 A from the solute) filled with TIP3P water. Na+ and Cl-counter ions were added to reach net-neutralization with salt excess to reach 150 mM NaCl. FF14SB and GLYCAM-06j force fields were used for protein and glycan residues, respectively. All MD simulations were performed using the Amber20 suite. The system was minimized by 2500 steps of steepest descent minimization and 2500 steps of conjugate gradient. Equilibration was performed using the NVT ensemble (200 ps) followed by NPT ensemble (200 ps), followed by the harmonic restraints in solute atoms. A last NPT equilibration step without restraints was performed for 500 ps. For each glycoprotein system and the IGHA1 system without glycans, the production run was performed at 298 K for 100 ns with a time step of 2 fs in triplicate. Hydrogen-containing bonds were constrained using SHAKE (Elber *et al*, 2011). Long-range electrostatic interactions were calculated using particle mesh Ewald, and short-range nonbonded interactions were calculated with a 9 A cutoff.

To analyze whether the glycans could affect the IGHA1 structure, the distance between each IGHA1 residue (only considering Cα) along the simulation trajectory and its position in the reference structure was computed. The first frame of the IGHA1 simulation trajectory without glycan (WT) was used as a structural reference. Then, the median of the average deviations in each triplicate in the simulations with glycan was subtracted from the median of the deviation without glycans (WT). The deviations were computed considering only the last half of the trajectory. All analyses were performed using VMD(Humphrey *et al*, 1996) and the R package Bio3D (Grant *et al*, 2021).

### Data visualization and processing

Heatmap diagrams, bar plots, and regression model plots were created using data visualization in the R (v3.6.0) environment and were also based on identified modified peptide sequences. The mechanistic hypothesis for OSCC diagnosis and metastasis in the salivary environment was visualized using BioRender (BioRender.com).

## Lead Contact

Further information and requests for resources and reagents may be directed to and will be fulfilled by Adriana Franco Paes Leme.

## Materials Availability

This study did not generate new unique reagents.

## Acknowledgments

We thank the contribution and support given by the Mass Spectrometry (MAS) Laboratory at Brazilian Biosciences National Laboratory.

## Funding and additional information

Funding was provided by FAPESP Grants (2009/54067-3; 2010/19278-0, 2016/07846-0; 2014/23888-0 and 2018/18496-6 to AFPL; 2014/06485-9; 2018/15535-0 and 2019/18751-9 to DCG; 2019/09692-9 to AGCN; 2019/21815-9 to AFBL; 2018/02180-0 to CMC; 2018/12194-8 to LDT) and CNPq Grants (470549/2011-4, 305851/2017-9 305748/2017-3, 428527/2018-3, 306279/2020-7, and 310392/2021-7 to AFPL).

## Author contributions

D.C.G. and A.F.P.L. designed research; A.R.S.S., M.A.L., T.B.B., A.C.P.R., collected, processed, and stratified clinical samples; D.C.G., S.Y., and A.G.C.N were responsible for saliva collection, performed LiP-MS sample preparation and LiP-MS screens; B.P. and R.R.D. performed LC-MS/MS data acquisition and mass spectrometer maintenance; D.C.G., L.D.T., C.M.C., A.F.B.L., G.P.T., H.V.R.F., F.M.S.P., G.A.C analyzed and interpreted data and led bioinformatics analysis across the project; D.C.G., A.G.C.N., L.D.T., C.M.C, A.F.B. L. and A.F.P.L. wrote the manuscript with contribution from all authors. D.C.G. and A.F.P.L. edited the manuscript and coordinated revision.

## This paper explained

### Problems

OSCC presents a low rate of survival, and patients also have a high rate of relapse and a low response to coadjuvant therapies. Therefore, the pursuit of identifying targets that can guide OSCC prognosis or diagnosis is important to progress in early diagnosis and better stratification of patients.

### Results

More proteins and pathways are brought up with LiP analysis associated with OSCC, TKT, and IGHA1 are both factors that show changes in protein conformation, probably through protein-protein networks or glycan composition exclusivity.

### Impact

Our findings suggest proteins with conformational changes in saliva associated with OSCC clinical features and different PTMs that can guide the search for sites with loss or gain of function as well as inquiring signaling pathways that would not be inquired about without structural accession.

## Data availability

The mass spectrometry proteomic data have been deposited in the ProteomeXchange Consortium (http://proteomecentral.proteomexchange.org) via the PRIDE partner repository (Vizcaíno *et al*, 2013) with the dataset identifier PXD042574. All other data supporting the findings of this study are available from the corresponding author on request.

## Supplemental Information

Twenty-five supplementary tables and three supplementary figures are provided within the manuscript.

## Abbreviations and nomenclature

OSCC: Oral Squamous Cell Carcinoma
N0: Absence of lymph node metastasis
N+: Presence of lymph node metastasis
TNM: Tumor (T), Nodes (N), Metastasis (M) classification system
LC-MS/MS: Liquid Chromatography-Tandem Mass Spectrometry
FDR: False Discovery Rate
LiP: Limited proteolysis
LiP-MS: Limited proteolysis-mass spectrometry analysis
DDA: Data dependent analysis.

