## Supplementary Figures for "Conformational changes in saliva proteome guides discovery of cancer aggressiveness related markers"

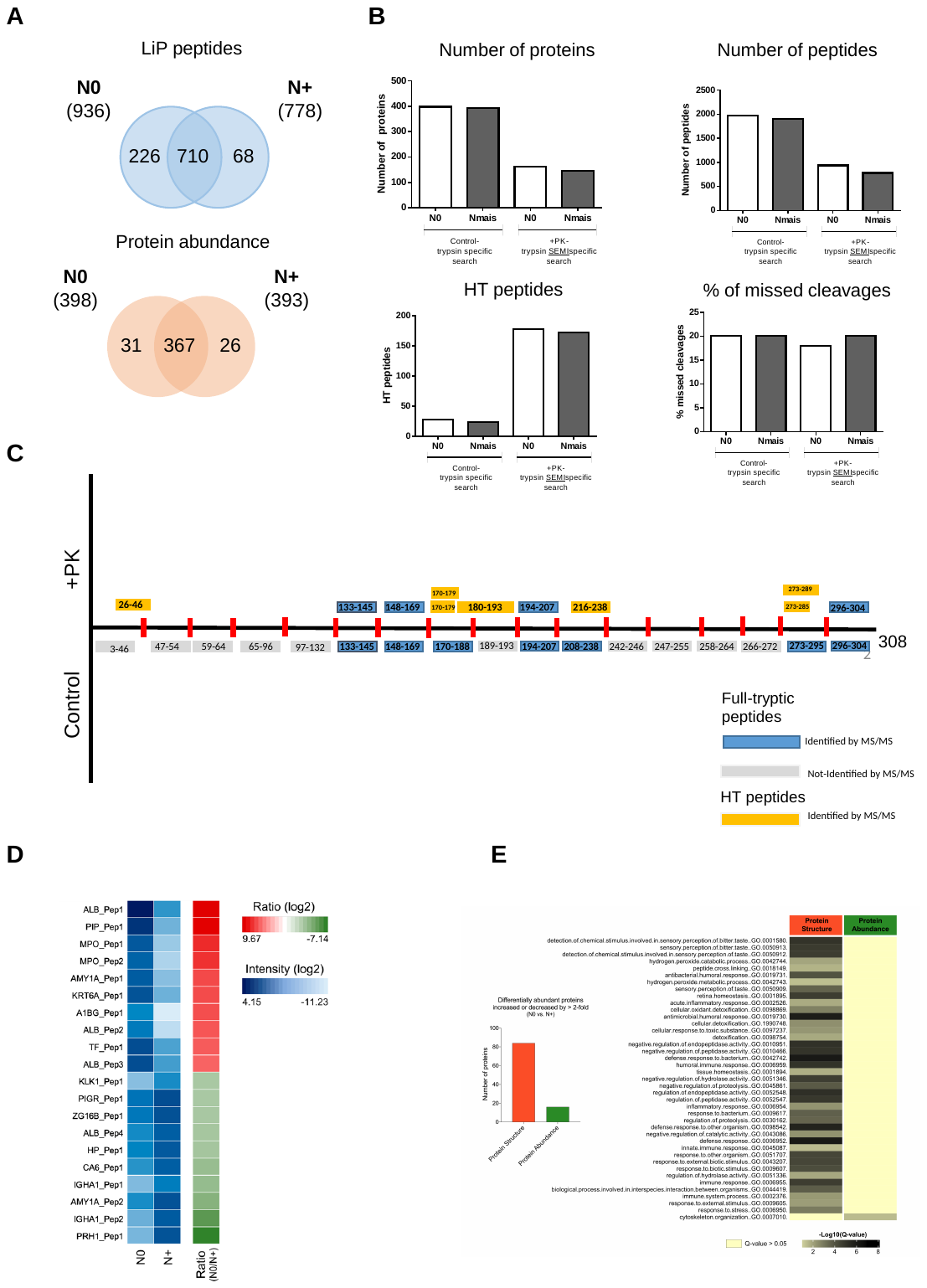

**Supplementary Figure 1.** LiP-MS optimization in saliva pooled samples (N0 and N+ patients). Relevance of structures of saliva in OSCC diagnosis and prognosis. (a) Venn diagrams illustrating the intersection of proteins and peptides that represent OSCC metastasis using “Database only search”. (b) The total number of proteins, peptides, half-triptic peptides (HT) and the percentage of missed cleavages out of the total number of peptides identified are reported for the different treatments (control vs PK). (c) As an example of a protein with conformotypic peptides accessible to PK treatment, the CA6 (Carbonic anhydrase 6) protein is shown represented by a black line; red boxes indicate the position of K and R residues. Blue and Orange boxes represent fully tryptic and half-tryptic peptides identified during the LiP-MS analysis, respectively. The position of the boxes reflects the position of the associated peptide along the protein sequence. (d) Heatmap presenting the abundance of peptides associated with OSCC metastatic phenotype (N+ or N0; FC≥2). Top-10 up and downregulated proteins showing intensity and their respective FC values. (e) Combined view of GO biological processes enriched for OSCC-metastatic associated proteins identified by LiP-MS/MS vs. abundance-based proteomics. GO biological processes were enriched using the Panther tool (FDR ≤ 0.01) and overrepresented GO processes were summarized by removing redundant terms in REVIGO. d) Protein abundance and structural changes were monitored with LiP-MS and compared between N+ and N0 with log2FC>2 ou <-2 (no statistical analysis was applied in the optimization assay). As a result, 16 and 84 proteins changed in abundance and structure, respectively. Heat map of GO biological processes were enriched using PantherDB ([http://pantherdb.org](http://pantherdb.org/)) considering the top-20 altered proteins in pooled saliva samples from N0 and N+ OSCC patients. Only significant biological processes using Fisher exact test and Bonferroni correction are shown. In total, 40 different biological processes were enriched considering structural readout against one process enriched based on changes in abundance readout. Illustrated in yellow are non-enriched processes in one or the other condition (Q > 0.05).

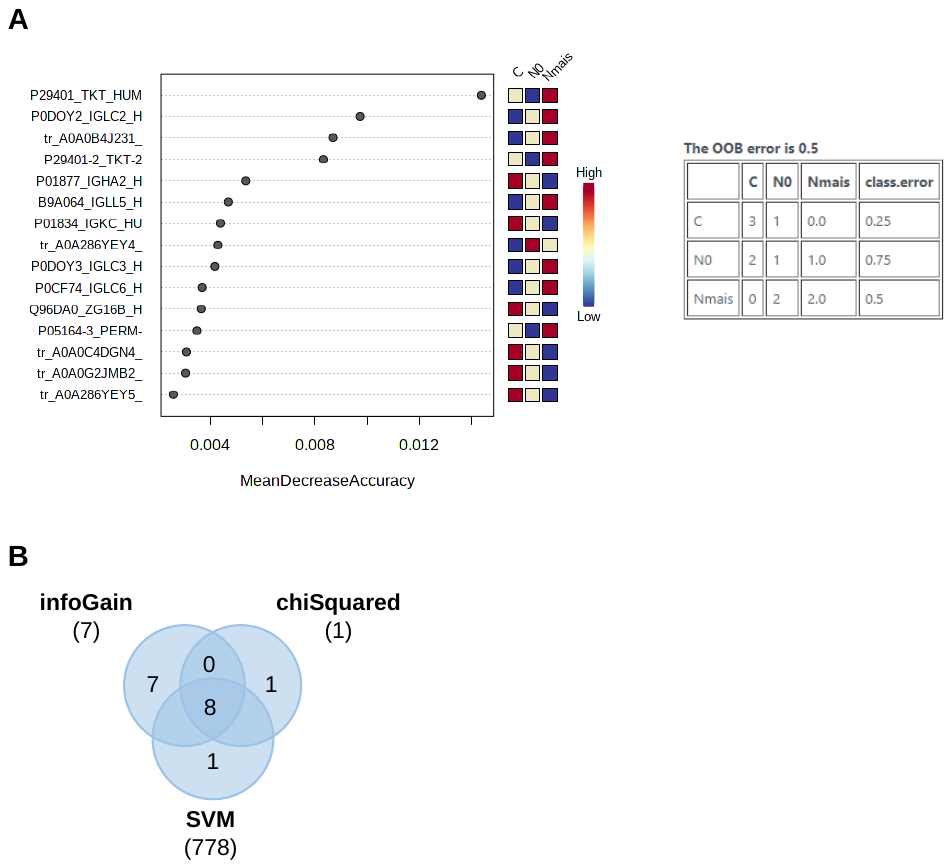

**Supplementary Figure 2.** (a) Variable importance plot (VIP) and confusion matrix from Random Forest analysis for prioritization of TKT prognostic protein. (b) Comparison of the common proteins in the three different ranking of attributes methods applied to the dataset. TKT peptides are among the eight common peptides.

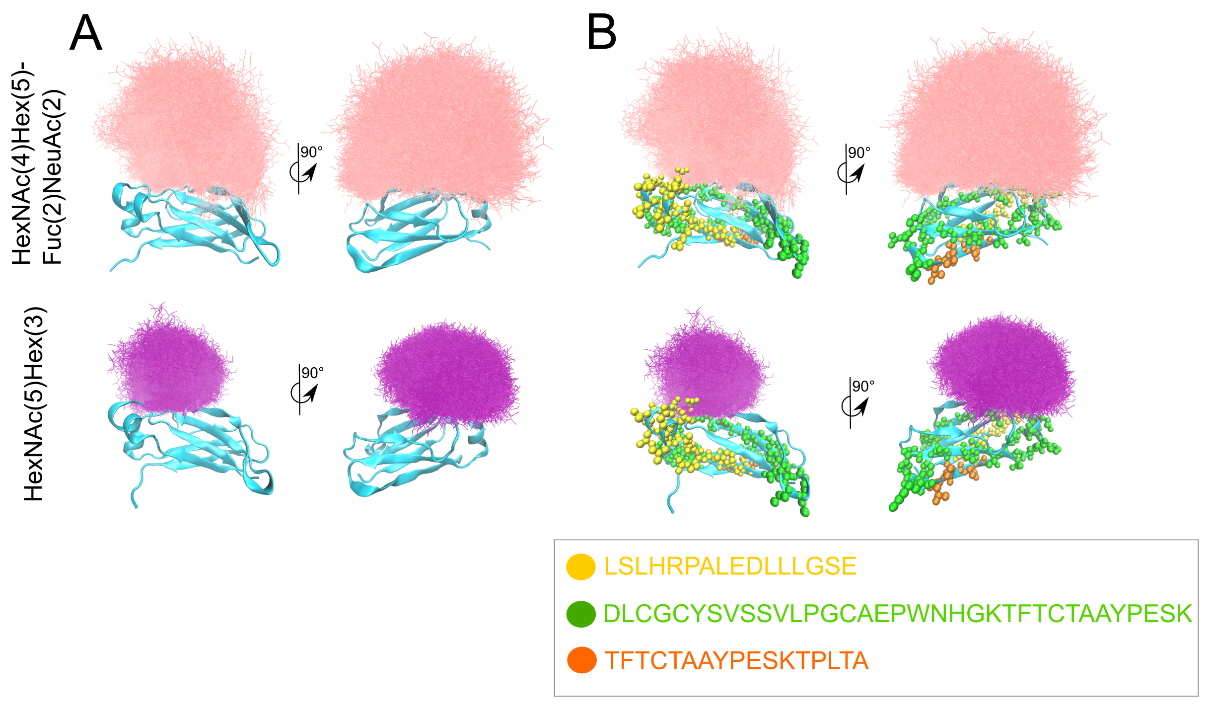

**Supplementary Figure 3.** Molecular dynamics simulations of IGHA1 Fc domain bound to the glycans HexNAc(5)Hex(3) and HexNAc(4)Hex(5)Fuc(2)NeuAc(2). (a) Superposition of the simulation trajectory aligned by the IGHA1 domain (in cyan) and presenting the glycan position (as lines) from every 0.2 ns of the 100 ns trajectory (in triplicate). Only the first frame of the IGHA1 is presented. Pink lines corresponds to HexNAc(4)Hex(5)Fuc(2)NeuAc(2) atoms and purple lines correspond to HexNAc(5)Hex(3) atoms. (b) Same as (a) but highlighting three conformotypic identified peptides as spheres. The sequence of each peptide and its color scheme are shown in the bottom box.
